# Asymmetric, dynamic adaptation in prefrontal cortex during dichotic listening tasks

**DOI:** 10.1101/683987

**Authors:** J. A. N. Fisher, I. Gumenchuk, O. Rogovin, A. G. Yodh, D. R. Busch

## Abstract

Speech comprehension relies on highly distributed, dynamically interconnected neuroanatomical loci. Accordingly, performance on complex speech processing tasks such as dichotic listening can be used to assess the integrity and health of many functional and structural aspects of the brain. Despite the potential merits as a clinical assessment tool, however, the neural substrates activated during dichotic listening remain relatively opaque at higher processing levels. Ultimately, this knowledge gap limits diagnostic use of the task. At the level of the prefrontal cortex, dichotic listening induces an asymmetric response wherein regions on the right hemisphere exhibit a higher functional activation than on the left. Superficially, this finding is counterintuitive given the left hemisphere’s dominance for speech and language. To obtain a more in-depth perspective on the potentially distinct roles of the right and left prefrontal cortex, we optically monitored cerebral blood flow in the dorsolateral prefrontal cortex (DLPFC) during dichotic listening tasks in human subjects. The method permitted us to avoid systematic experimental confounds that functional magnetic resonance imaging (fMRI) measurements suffer from, namely the influence of scanner noise. In addition to reproducing the documented larger activation amplitude in the right hemisphere, we also found that repeated listening task blocks were associated with altered kinetics of blood flow in the right, but not the left DLPFC. Interestingly, subjects with the most prominent regional blood flow changes in the right hemisphere also displayed large distortion product otoacoustic emissions (DPOAEs) in the left ear, possibly signaling a correlation between prefrontal activity and top-down listening control infrastructure through medial olivocochlear efferent projections to the inner ear. Overall, our results suggest that the right prefrontal cortical regions play an active role in optimizing task performance.

## Introduction

Attending to and comprehending speech leverages a highly distributed network of functional neural infrastructure. Unsurprisingly, damage to or dysfunction of any aspect of the central nervous system often interferes with aspects of speech perception, such as comprehension in noisy environments (Musiek et al., 2004; Lew et al., 2007). Behavioral tasks that probe speech processing therefore hold potential as more general neurological assessment instruments. From a practical perspective, they are attractive because they are easy to administer and there are well-established normative values. Among such tests, dichotic listening tasks (Kimura, 1961, 1967), which assess an individual’s ability to retain or selectively attend to simultaneously presented sounds that differ between the ears, have been extensively explored and can uncover otherwise occult neural dysfunctions for a very wide range of conditions (Hugdahl, 2003). Despite this promise, the links between performance on dichotic tasks and underlying neural mechanisms are not well understood. Given that dichotic listening tests are highly sensitive to brain health, understanding the underlying neural substrates for this capability could offer quantitative metrics that are even more sensitive to otherwise undetectable conditions.

One of the most robust empirical characteristics of dichotic listening tests is that right-handed subjects consistently retain and repeat information presented to the right ear more accurately than the left, a phenomenon termed a “right ear advantage” (REA) (Berlin et al., 1973; Bryden et al., 1983). Mechanistic explanations for the REA center around the fact that left hemisphere auditory areas are dominant for language processing and that retaining info from the left ear is an intrinsically “uphill” task. Imaging results using ^15^O positron emission tomography (PET) (O’Leary, Hugdhal 1999, Hugdahl 2000) indeed demonstrate that at temporal cortical auditory areas, there is a left-biased asymmetric activation during dichotic listening. Otherwise REA has been proposed to reflect an embedded anatomical bias for information transfer, specifically through the corpus callosum (Kimura, 1967; Zaidel, 1983; Westerhausen and Hugdahl, 2008) or else an active left-hemisphere priming due to attentional networks (Kinsbourne, 1970), potentially of subcortical origin (Kinsbourne, 2003; Zaidel and Iacoboni, 2003). In fact, “split-brain” patients who have had their corpus callosum transected are not able to recall numbers presented to the left ear (Milner et al., 1968). These same active mechanisms have been proposed to underlie the fact that people still perform reasonably well on left ear reporting (Strouse and Wilson, 1999) as well as the fact that people can selectively switch attention and enhance recall from ear to ear (Kompus et al., 2012).

At the level of prefrontal cortex, however, the left hemisphere dominance during dichotic listening tasks is not as apparent. In fact, imaging studies have found that dichotic listening tests can evoke comparatively larger activation in the right prefrontal cortex (Pugh et al., 1996; Larisch et al., 1999; Jancke and Shah, 2002). Multiple rationales have been proposed for these findings, such as a necessity of the right hemisphere to devote additional compensatory activation to retain info from the left ear in order to overcome the hemisphere’s relative disadvantage for speech and language (Kompus et al., 2012). Alternatively, the same callosal or subcortical substrates proposed to underlie REA may be sufficiently multifunctional to evoke this greater activation (Jäncke and Steinmetz, 1994). The right PFC activity has also been proposed to reflect a more generalized, modality-neutral attentional network (Hugdahl et al., 2000). Of course, given the qualitative nature of all of these models, these mechanisms are not mutually exclusive (Jancke and Shah, 2002).

Given the potential relevance of right PFC to more general, modality-neutral performance optimization, one approach for assessing this possibility would be to monitor the timecourse of regional functional activation in PFC during repeated behavioral tasks. Repetition on any behavioral task alters performance, an outcome that involves dynamic plasticity in myriad cognitive processes such as working memory management, attention, and information processing. Even at the level of concise sensory-evoked responses, repeated stimuli elicit progressively smaller responses only in regions closely involved with execution of that task or handling of that information (Grill-Spector and Malach, 2001; Epstein et al., 2008). Regions closely involved in orchestrating this process, putatively right PFC, would be expected to exhibit dynamic activation that co-varies with changes in task execution and performance.

Dynamic aspects of functional activation during dichotic listening, however, have been challenging to acquire at high temporal resolution using magnetic resonance imaging (MRI)-based modalities because of technical limitations related to scanner noise. For example, in their functional MRI (fMRI) study on dichotic listening, Jancke and Shah (2002) found that the noise level was 70-80 dB SPL even after attenuation by protective headphones. Facing the same limitations, Thomsen et al. (2004) identified activated regions but were limited in their ability to collect multiple time points during listening task blocks. In a related study by Schmithorst et al. (2013), the signal-to-noise of fMRI measurements was low enough that sophisticated statistical methods were required in order to attribute regional activity to the task. Finally, scanner noise can alter the auditory pathway itself by eliciting reflexive middle-ear muscle contractions (stapedius reflex), or by activating the medial olivocochlear efferent reflex, which attenuates afferent auditory signals in response to loud sounds. Although a popular strategy is therefore to scan in between listening trials to avoid the confound, these efferent effects (which are activated at the intensities involved in MRI scanning) can persist up to 50 sec after loud sounds (Sridhar et al., 1995). In terms of other clinical modalities, PET has insufficient temporal resolution, and non-imaging modalities such as EEG or magnetoencephalography (MEG) do not have the requisite spatial resolution to probe regional changes in activation.

To avoid these systematic confounds, we monitored regional cerebral blood flow (CBF) during a repeated, free-recall dichotic digits listening task in heathy subjects using an optical technique, diffuse correlation spectroscopy (DCS). DCS takes advantage of the dynamic scattering of light from moving red blood cells to directly measure cerebral blood flow and is particularly sensitive to flow in the cortical microvasculature due to the high absorption (and thus low probability of photon escape) in larger blood vessels (Durduran and Yodh, 2014). We monitored changes in acoustic properties of recorded verbal responses as a proxy for performance changes. Additionally, to explore whether PFC functional asymmetry may be relevant to top-down modulatory mechanisms, we measured distortion product otoacoustic emissions (DPOAEs), whose magnitude has been found to vary with auditory and spatial attention (Andéol et al., 2011a; Wittekindt et al., 2014).

## Materials and Methods

### Participants

Eight healthy, right-handed subjects (5 women, 3 men age range 16-37) participated in this study, which was approved by the Institutional Review Board at New York Medical College and at Westchester Medical Center. Informed consent was obtained from the subjects prior to the measurement session, and subjects were asked to refrain from consuming caffeinated beverages the day of the experiment.

Subjects were asked to confirm that they could clearly and comfortably hear the sound stimuli in the study room with all equipment and cooling apparatus active.

### Listening Tasks

Subjects wore insert earphones (ER3-14A, Etymotic Research, Elk Grove Village, IL) and performed diotic and dichotic digits listening tasks. During dichotic digits task blocks (Fig. 1), each ear was presented with three sequentially spoken numbers selected randomly from one to ten, but excluding “seven” so that all words had only one syllable. Each ear was presented with three different, non-repeating numbers so that in total, a subject was presented with six different, randomly selected numbers. Spoken digits were presented as simultaneous pairs, one digit per ear, at a pace of roughly 2 words / sec. For example, a subject might hear “two, five, four” in one ear while simultaneously hearing “eight, ten, one” in the other. Subjects were asked to verbally report the six numbers they heard, regardless of ear or order (i.e. free recall). Subjects were advised that if they could not remember one or more of the numbers, they should just guess. Subjects were asked to say the numbers clearly, yet at a normal speaking volume, and were informed that their responses were being recorded.

**Fig. 1:**
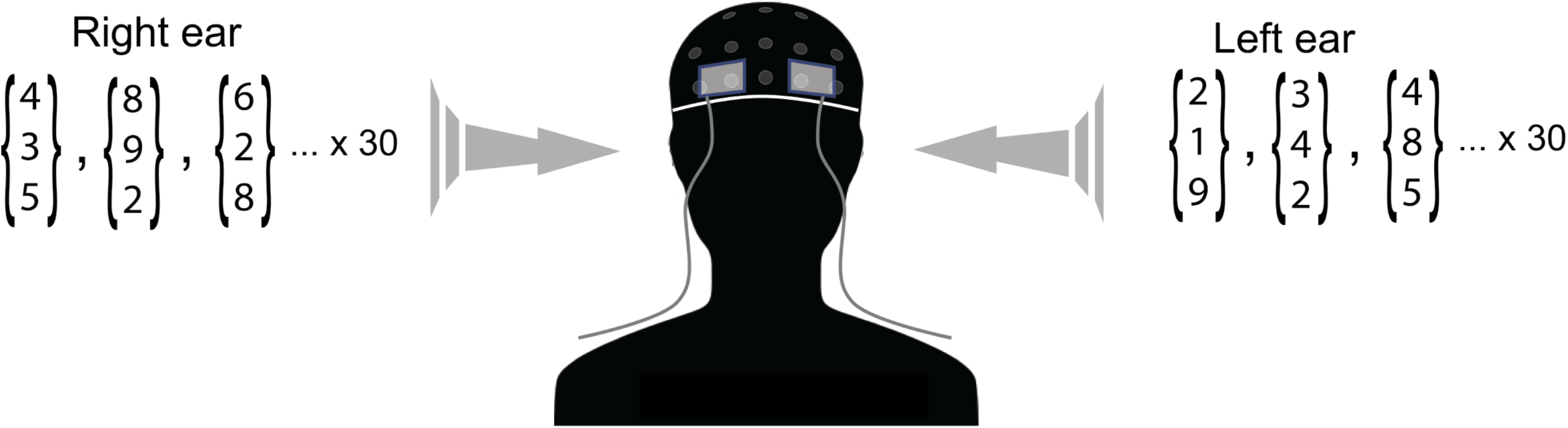
Illustration of experimental protocol. Optical probes embedded in flexible elastomer were placed at locations AF7 and AF8 (10-10 international electrode placement system) and secured under a neoprene EEG cap. The three-digit arrays depict the groups of three spoken digits that were selected randomly and delivered to each ear through calibrated insert earphones during dichotic listening tasks.

During diotic digits task blocks, both ears were simultaneously presented with the same numbers in unison from a single randomly selected set of three numbers so that in total only three unique numbers were heard. Speaking can introduce mechanical artifacts in both CBF and EEG measurements. To ensure that there was not an uneven influence of mechanical artifact on diotic vs. dichotic listening tests, subjects were instructed during the diotic tests to twice recite the three numbers they heard, e.g. “five one two, five one two.”

Spoken numbers were selected and stimulus waveforms assembled using custom LabVIEW code (National Instruments, Austin, TX) and output as analog waveforms through a USB-based DAQ unit (USB-6215, National Instruments). The voices used for constructing the stimulus were computer-generated male voices. Analog waveforms were amplified by a stereo power amplifier (SA1, Tucker-Davis Technologies, Alachua, FL) and input to earphone amplifiers (ER-1, Etymotic Research, Inc.) that were coupled to insert earphones through plastic tubing. The average output at the earphones was slightly adjusted between subjects but was within 65 ± 5 dB SPL.

### Study Design

Experimental sessions lasted approximately one hour. After an initial consenting process and brief interview, during which we described the breakdown and flow of the experiment, the subject’s DPOAEs were measured in the office. Subjects were then moved to an acoustically isolated room and remained there for the rest of the experiment. After fitting the subject with an EEG cap, gelling electrodes and positioning optical probes, subjects were instructed how to perform the diotic and dichotic listening tests. Subjects were also given a sample of what each one would sound like, and how to properly respond (e.g. announce the numbers in an even, clear voice).

The experiment followed a block paradigm, and consisted of the following phases: (1) initial rest (2 min), diotic digits (∼4-5 min), rest 2 (2 min), dichotic digits block 1 (∼4-5 min), rest 3 (2 min), dichotic digits block 2 (∼4-5 min), rest 4 (2 min). Subjects were informed when blocks were starting and when they had ended. There was typically 10-15 seconds delay between the end of one block and the beginning of the next, just to give subjects a moment to slightly adjust seating position if they were experiencing discomfort. During the experiment, the room was nearly entirely dark, except for computer monitors which faced away from the subject. Subjects were instructed to keep their eyes open (with the exception of blinking) and to try to maintain their gaze fixated on a target (black cross on white background) ∼8 feet from their face.

### Measurement of Cerebral Blood Flow Using Diffuse Correlation Spectroscopy

Regional, cerebral blood flow (CBF) was measured using diffuse correlation spectroscopy (DCS), which is a near-infrared interferometric technique that directly measures microvascular blood flow (Durduran et al., 2010a; Buckley et al., 2014). DCS utilizes the interference pattern formed on tissue surface following illumination by a long-coherence length laser. Fluctuations in the interference pattern are related to the displacement of red blood cells and can be utilized to compute a blood flow index. Changes in this index from baseline reflect changes in blood flow (Mesquita et al., 2013; Shang et al., 2017), as validated by multiple studies against Doppler ultrasound (Buckley et al., 2012), fluorescent microspheres in piglets (Zhou et al., 2009), and MRI techniques (i.e., arterial spin-labeled perfusion, phase-encoded velocity mapping) (Durduran et al., 2010b; Jain et al., 2014). This study utilized a high-speed variant of DCS, capable of measurement rates of up to 50 Hz (Wang et al., 2016). Briefly, continuous wave, long coherence length lasers (785 nm; 80 mW; DL785-100-3O, 830 nm; DL830-100-3O, CrystaLaser Inc., Reno, NV) were used to illuminate the scalp via a prism-coupled multimode fiber (200 μm diameter, OZ Optics, Ottawa, Canada). Remitted light that travelled through the head was detected by prism-coupled single mode fibers (780HP, 6 μm core diameter, Fiberoptic Systems Inc., Simi Valley, CA; Nefern, East Granby, CT) located 2.5 cm from the source. Each detector fiber directs light to a single photon-counting APD (i.e., each fiber-detector combination is independent, SPCM-AQ4C, Excelitas, Quebec, Canada). Correlation functions derived from co-located detectors were averaged. Prisms coupled to source and detector fibers were embedded in flexible pads made from pourable elastomer that were positioned on the scalp at locations corresponding to AF7 (left) and AF8 (right) in the 10-10 EEG electrode placement system (Chatrian et al., 1985). The pads were slipped under a neoprene EEG cap (Enobio, Neuroelectrics, Inc., Cambridge, MA), which held them firmly, yet comfortably in place. Baseline optical properties were held fixed in the analysis (reduced scattering coefficient 10/cm; absorption coefficient 0.1/cm). Potential errors in the blood flow index (BFI) calculation due to a mismatch between the assumed and actual optical properties are minimized by utilizing the ratio between the BFI at each time point and that from the baseline period.

### EEG Measurements

EEG was recorded wirelessly using the 10-10 international electrode placement system (Enobio 20, Neuroelectrics). The headcap was fitted with wet electrodes (NE032, Neuroelectrics) which were then filled with Sigma Gel (Parker Laboratories, Fairfield, NJ). Channels with an impedance less than 10 kΩ were included in the analysis for this paper. Analog signals were sampled at 500 Hz and transmitted to a PC via Bluetooth. Data was analyzed offline using custom Matlab (Mathworks, Natick, MA) functions as well as EEGLab (Delorme and Makeig, 2004). Raw EEG data was bandpass filtered from 2-100 Hz.

### Distortion Product Otoacoustic Emissions (DPOAEs)

DPOAEs were measured using an Otodynamics Echoport 292 clinical binaural diagnostic device using ILO 6 data acquisition and analysis software. Emissions were elicited by delivering to the ear two simultaneously presented pure-tone frequencies, *f*_1_ and *f*_2_ that varied in frequency, yet were maintained at a ratio of *f*_2_/*f*_1_ = 1.22. Tones were delivered through ear probes that had two separate transducer elements as well as a microphone tube, all integrated into a single foam tip inserted into the ear canal. Due to the active, nonlinear mechanical properties of cochlear hair cells, these pure tones elicited emissions at other frequencies including a prominent emission at 2*f*_2_-*f*_1_. The sound pressure level (dB SPL) of the emitted sounds recorded within the ear canal at 2*f*_2_-*f*_1_ was recorded for 2*f*_2_-*f*_1_ at frequencies 1, 1.5, 2, 3, 4, and 6 kHz.

### Analysis of Vocalizations

Subjects’ verbal responses were recorded with a mono microphone (ACM 1b, Cyber Acoustics, Vancouver, WA). Acoustic artifacts and experimental pauses such as the silent periods between trials and rest periods were removed from the waveform using Audacity software (Audacity Team). The data was subsequently analyzed using custom Matlab functions. We quantified the duration of responses as the time elapsed between the first and last reported numbers. Specifically, vocal response duration was measured from the first to the last point in time of each trial where the absolute magnitude of the recorded waveform exceeded the RMS ambient room noise level.

## Results

### Auditory Task Performance

Overall, subjects performed well on the dichotic digits task. In the first block, left and right ear accuracies summed over all subjects were 87% ± 3% and 95% ± 1% respectively (mean ± standard error of the mean). In the second block, accuracies for left and right ears were 90% ± 3.7% and 94% ± 1.5%, respectively. For each ear, these values were not statistically different between blocks. These values, including the left/right asymmetries, are consistent with normative values for three-digit dichotic listening tests in the same age group (Strouse and Wilson, 1999). After experimental sessions, when asked about their general comfort level, all participants felt that the second dichotic listening block was “easier.” Most participants noted that as they progressed through the dichotic digits task blocks, they adopted the strategy of focusing on this information from one ear while relying on “passive” recall to retain and report the digits delivered to the other ear.

### Cerebral Blood Flow

As depicted in Fig. 2A and B, listening tasks induced functional changes in CBF at recording locations AF7 (left) and AF8 (right) in the 10-10 EEG electrode placement system (Chatrian et al., 1985), corresponding to the left and right DLPFC (Brodmann areas 46L and R). As summarized in Fig. 3, averaged over all subjects, both diotic and dichotic digits tasks evoked statistically significant changes in CBF relative to the preceding rest phase. For all blocks, evoked changes in CBF were of greater magnitude in the right hemisphere than the left. Additionally, dichotic tasks evoked larger functional changes than the diotic task. The diotic digits task evoked a relative ΔCBF of 22% ± 6% and 12% ± 2% in the right and left dorsolateral prefrontal cortex, respectively. The subsequent dichotic task block evoked changes of 35% ± 8% (right) and 19% ± 3% (left) for the first block. In the second dichotic block, the average CBF increased further (41% ± 9%) and the increase was statistically significant compared with the first block (*P* = 0.014, Mann-Whitney U test). In contrast, there was no statistically significant change in left hemisphere signal compared with the prior dichotic block.

**Fig. 2:**
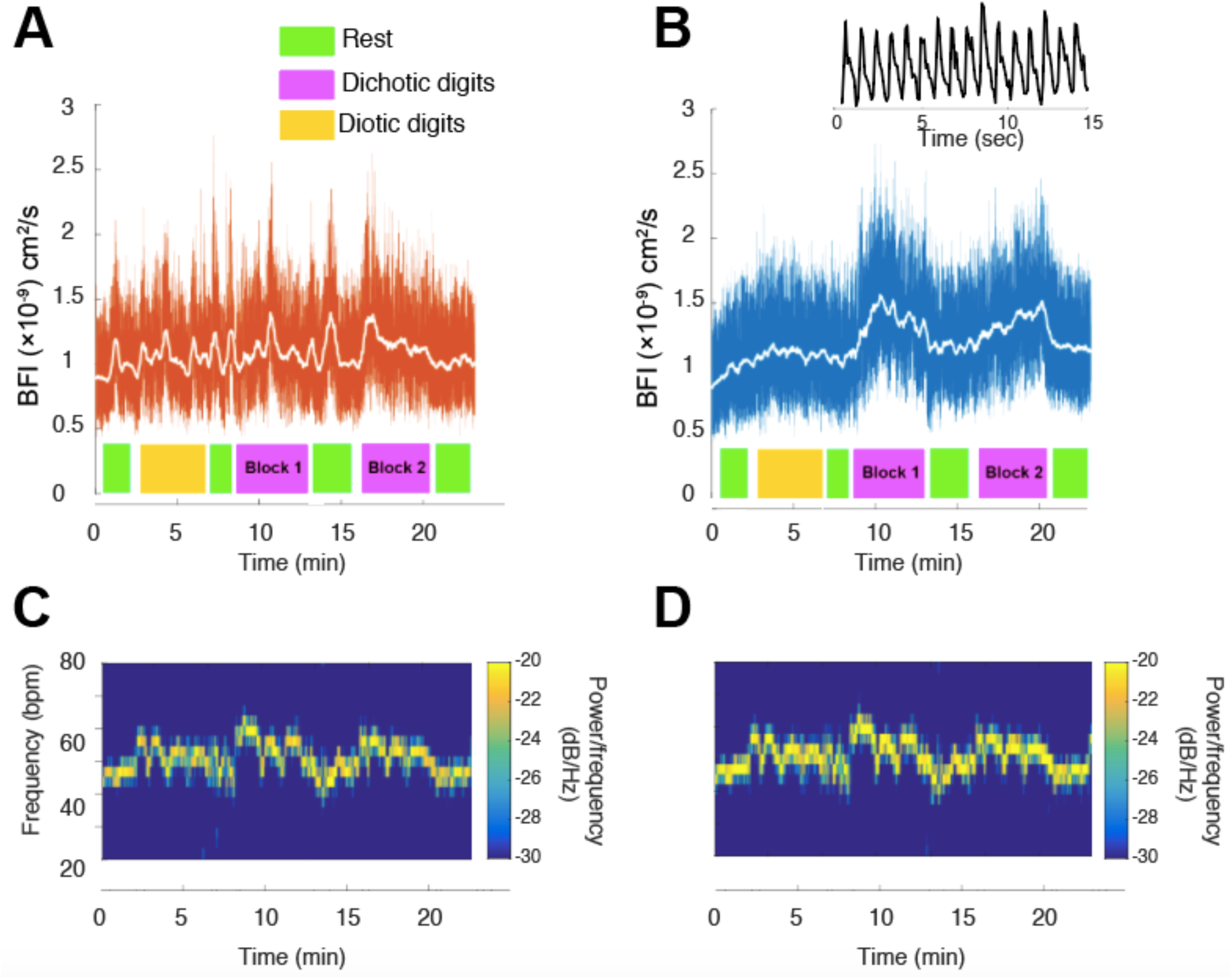
Cerebral blood flow (CBF) in prefrontal cortex during listening tasks in one subject. (A) and (B) depict relative cerebral blood flow measured optically at scalp positions corresponding to the left (A) and right (B) dorsolateral prefrontal cortex during a series of listening task blocks and rest periods. Values are displayed as blood flow index (BFI). The white trace superimposed on the data in (A) and (B) shows the panel’s same data smoothed with a 5-second moving average filter. Note that much of the ‘noise’ is in fact flow changes due to dynamic physiology (breathing, heart beat). The inset above (B) shows a 15-second snippet of the optical raw data showing pulsatile flow due to the cardiac cycle. Representative of the population averaged results, CBF at the recording locations increased most significantly during dichotic listening task blocks, and the signals were much higher on the right hemisphere. (C) and (D) depict spectrograms of the optical signals in (A) and (B), respectively, in the range of 20 – 80 Hz. These signals reflect heartrate during experiments and are symmetric, unlike the left-right asymmetry visible in the raw data plots in (A) and (B) above.

**Fig. 3:**
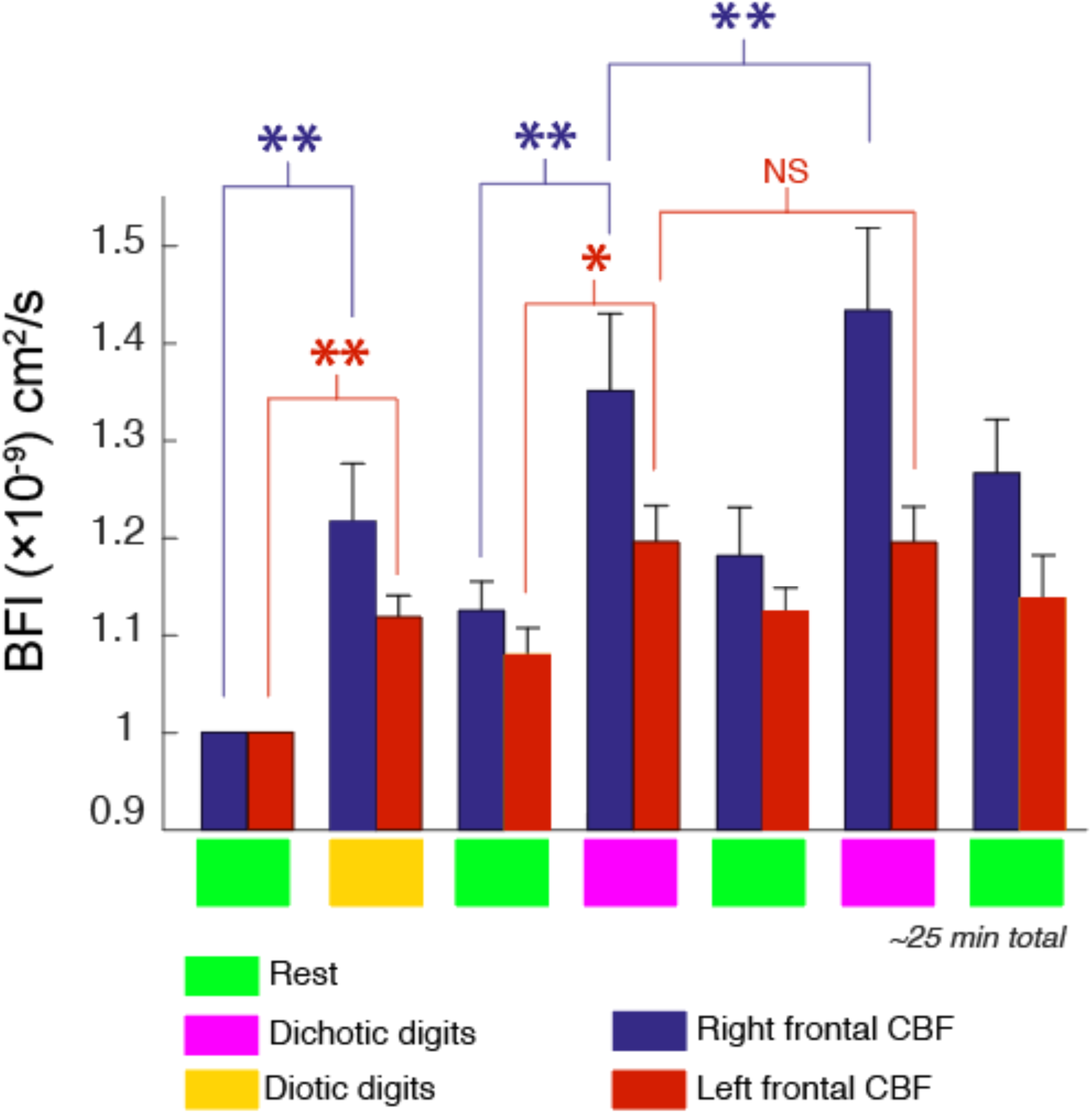
Cerebral blood flow changes during listening tasks averaged over all subjects. Blue and red bars depict relative changes in, respectively, right and left hemispheres. The values are normalized to the first rest period. Both diotic and dichotic tasks elicited significant increases in CBF. When comparing CBF changes during dichotic listening task blocks, there was a statistically significant amplitude increase from block 1 to block 2 in the right hemisphere. In the left hemisphere, however, there was no such statistically significant difference between the CBF responses in block 1 vs. block 2. Error bars represent standard error of the mean, and asterisks indicate statistical significance as: * = *P* <0.05, ** = *P* < 0.01.

In addition to magnitude changes, the kinetics of within-block ΔCBF changed as well from the first to the second dichotic listening tasks. Fig. 4 shows one representative subject’s CBF during the two dichotic digits tasks. Superimposed on the full data trace are local polynomial fits during the dichotic listening blocks, which we used to quantify the nonlinear temporal dynamics of the response. Compared with the first dichotic block, the functional ΔCBF in the right hemisphere exhibited a slower increase and reached a peak value on average 77 ± 4.3 s later than the first block. No statistically significant change in kinetics was noted in the left hemisphere, however. Subjects with low amplitude ΔCBF (3 out of 8) yielded very poor fits and were not included in this analysis; however, assessed among the remaining subjects, the kinetics change was still statistically significant (*P* = 0.028, Mann-Whitney U test).

**Fig. 4:**
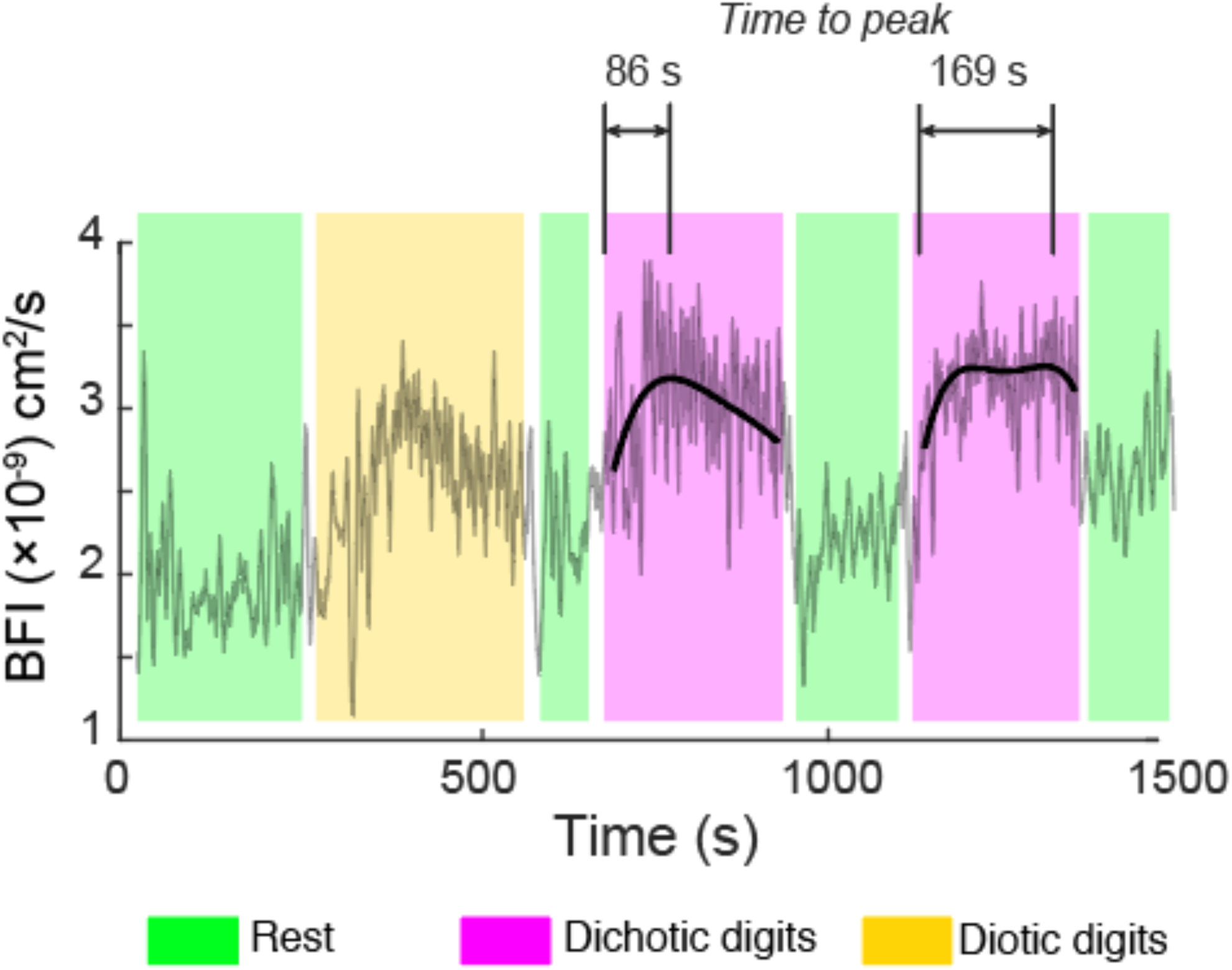
Activation kinetics in the right PFC differs between the two dichotic listening blocks. Here, local third-order polynomial fits are superimposed over the raw optical data during the dichotic listening periods. The shaded regions indicate rest (light green), diotic (light yellow), and dichotic (light purple) blocks. The increased time-to-peak in diotic block 2 that is highlighted here is representative of the population average. No statistically significant corresponding change in kinetics was found in the left hemisphere.

Although the measurements appeared to contain an abundance of random high frequency noise, as can be seen in the inset for Fig. 2B, the amplitude fluctuations largely reflect pulsations in blood flow due to the cardiac cycle. As shown in Fig. 2C and D, spectral analysis of the optical signal revealed that listening tasks transiently increased heartrate. However, the increases in heartrate were hemispherically symmetric and the slow temporal evolution of the 40-70 Hz frequency band spectral density did not match the corresponding slow trends in the unfiltered CBF data.

### EEG

During dichotic listening task blocks, spectral features of the ongoing EEG signal at frontal and parietal recording locations displayed changes that were statistically significant. As shown in Fig. 5A, the resting EEG spectrum featured prominent alpha (8-13 Hz) and beta (14-30 Hz) spectral features as well as a low gamma peak centered at ∼35 Hz. While the diotic digits task did not elicit significant changes in the EEG spectrum, dichotic task blocks were accompanied by significant changes in spectral power in the low-gamma range (30-50 Hz). Summed over all experiments, the trends are depicted in Fig. 5B. Whereas the CBF data for both hemispheres demonstrated a greater signal in the second dichotic block, the EEG amplitude was slightly, though not statistically significantly, greater for the first block.

**Fig. 5:**
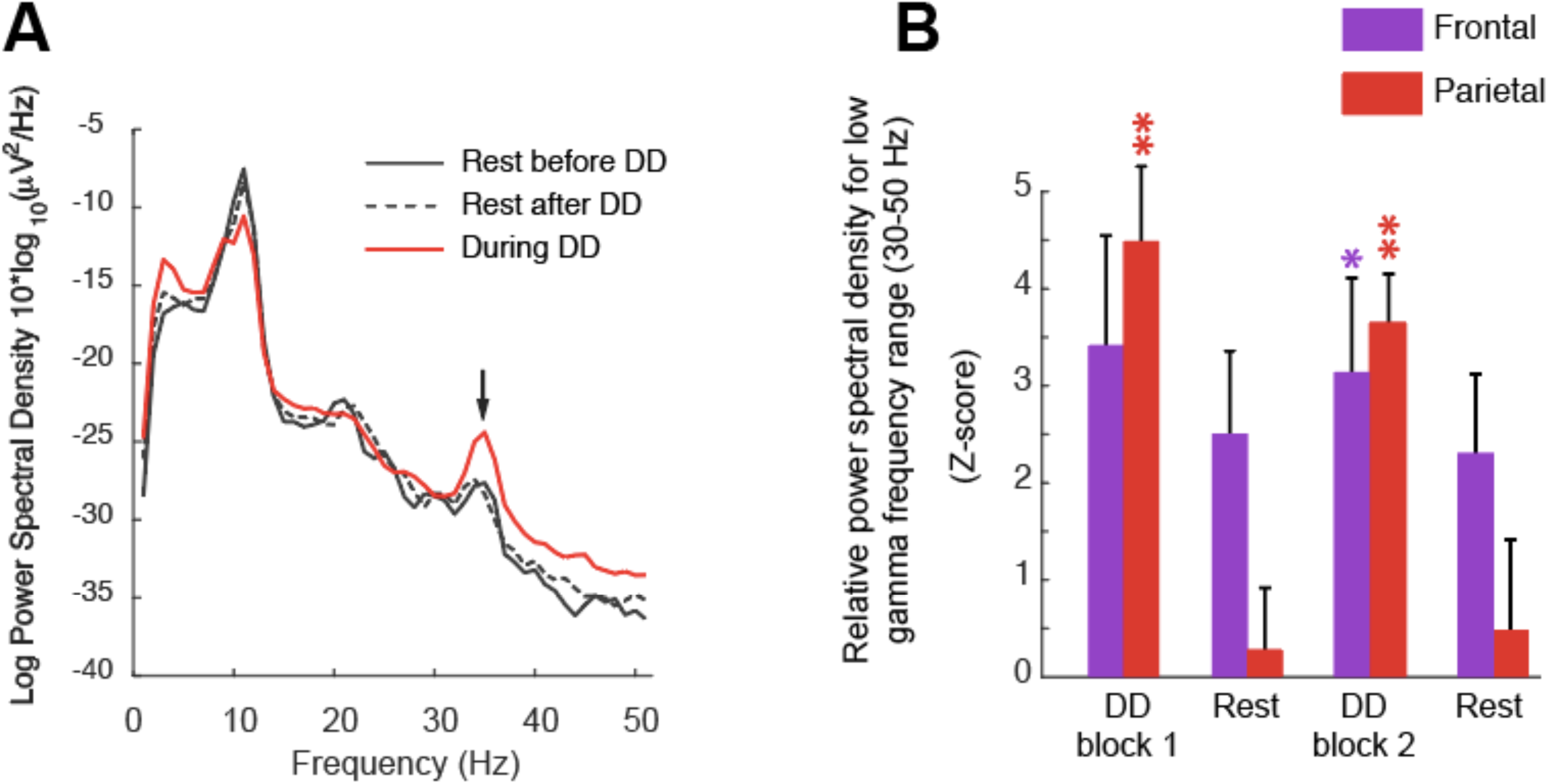
Dichotic listening increases EEG low gamma power in frontal and parietal recording locations. (A) EEG spectrum in a representative subject during dichotic listening periods (red trace) as well as rest periods before and after (solid and dotted black traces, respectively). The trace is averaged over parietal EEG recording locations. (B) Bar chart summarizing low-frequency gamma spectral power at frontal (purple) and parietal (red) recording locations during rest and dichotic listening periods. Because the baseline EEG power measurements varied between subjects, results are expressed as z-score, which represents the ratio of averaged spectral power during the measurement period (i.e. rest or task) to the standard deviation of pre-block fluctuations. Values with *z*-score > 2 were considered significant. Asterisks indicate degrees of statistical significance, as detailed in the caption for Fig. 3.

### Speech Patterns

While the most direct metric for performance enhancement would be changes in response accuracy, the baseline accuracy was so high that we could not detect a significant change in performance across subjects, despite the fact that all subjects reported the second block being subjectively less difficult. As a metric, albeit indirect, for subjective task difficulty, we therefore assessed the speech patterns of subjects during verbal responses. When subjects were unsure or hesitant during digit recall, they repeated the numbers more slowly. We quantified this effect by measuring the total duration between the first and last spoken words, specifically defined as the points at which the recorded sound exceeded the RMS noise level by two standard deviations (i.e. *z*-score > 2). Fig. 6A depicts a representative subject’s spectrogram of the recorded sound during verbal responses. Verbal responses consisted of six spoken numbers corresponding to the six digits total (three per ear) subjects were presented with. In the diotic listening task, when both ears were presented with the same three digits, the subject was asked to repeat the three numbers twice. Compared with the responses during the diotic digits task, the total span of the verbal responses increased during the first block of dichotic digits, however not the second block. These trends are quantified in Fig. 6B, in which durations are normalized to the averaged durations during diotic tasks. These values represent the average over all subjects.

**Fig. 6:**
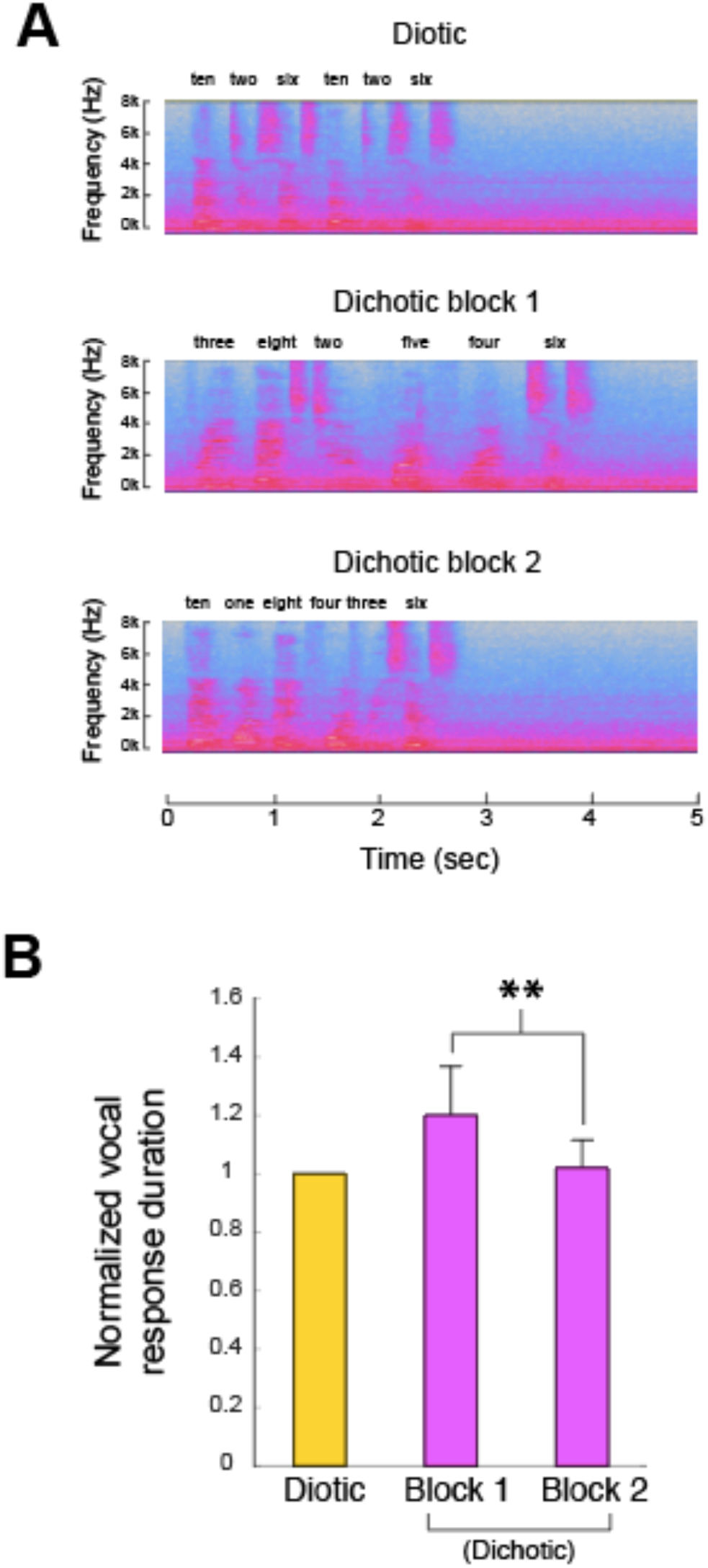
Analysis of verbal responses to listening tasks. (A) shows short segments of the spectrogram of a representative subject’s verbal responses. The panels illustrate how speaking rhythm can be visualized in the time-frequency plots. The prolonged verbal response duration is mostly evident in the first dichotic listening block. (B) Summarizes the average total duration of the free-recall responses following auditory stimuli. Vocal response duration was defined based on the first and last time points where the total recorded audio amplitude exceeded the maximum ambient noise level in the room that was present during that test. Shown here are the average durations of a vocal response for diotic and dichotic task blocks, all normalized to the diotic response durations. The reduction in response duration visible when comparing dichotic block 1 to block 2 was significant (*P* = 0.0079, Mann-Whitney U test).

### Correlation Between Cerebral Blood Flow and Distortion Product Otoacoustic Emissions (DPOAEs)

When subjects were examined individually, there appeared to be a nearly binary distinction in the magnitude of DPOAEs. Three out of the 8 total subjects demonstrated significantly lower DPOAE magnitudes for frequencies below 4 kHz (Fig. 7A-B), and the differences were more pronounced in the left ear. Interestingly, subjects with “low” vs. “high” DPOAE magnitudes were also quantitatively separable in CBF responses; subjects with high DPOAEs below 4 kHz in the left ear also demonstrated functional ΔCBF that had a *z*-score >2 for the first dichotic digits block, relative to noise levels in the preceding rest block (Fig. 7 C-D). Additionally, subjects with low left-ear DPOAEs appeared to have less ΔCBF asymmetry than subjects with high left-ear DPOAEs. Although the error magnitude was high due to small sample sizes, high DPOAE subjects had average right/left asymmetries of 13% ± 8% during the first dichotic digits block, whereas low DPOAE subjects had average asymmetry of 4% ± 9% during that same block.

**Fig. 7:**
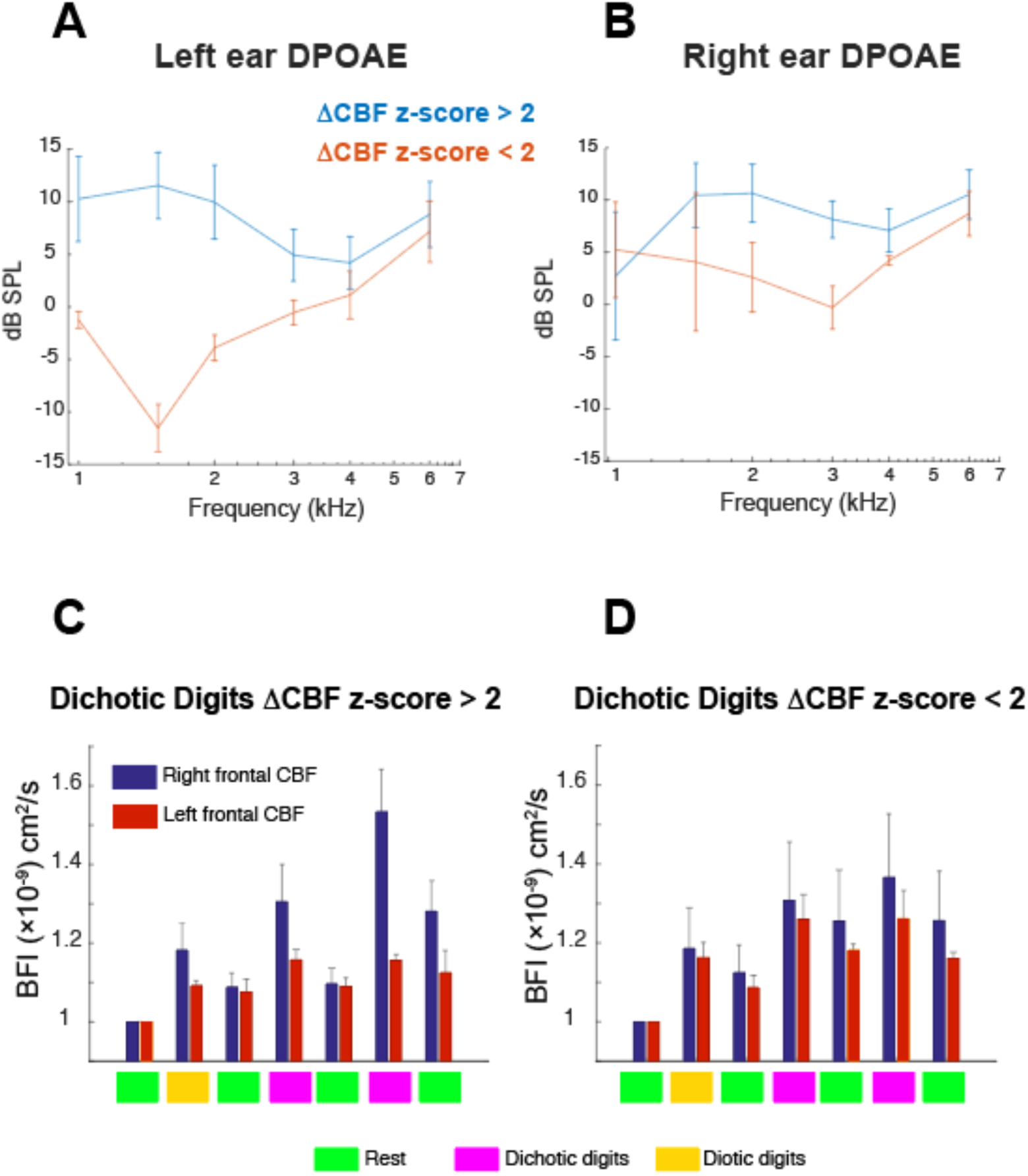
Distortion product otoacoustic emissions (DPOAEs) in subjects with low and high CBF changes during listening tasks. (A) and (B), respectively, show otoacoustic emissions recorded in the left and right ear of subjects segregated based on the magnitude of their CBF response during dichotic listening tasks. Bar charts in (C) and (D), respectively, depict the summed CBF trends in the populations of “high” and “low” response magnitude subjects. Specifically, the signal magnitude was evaluated based on the difference between dichotic block 2 and the preceding rest period.

## Discussion

Overall, by using optical techniques as an alternative to fMRI, we were able to temporally resolve the dynamics of an asymmetric activation in prefrontal cortex. The fact that these asymmetric hemodynamic trends are accompanied by indications of reduced task effort during dichotic listening suggests that right PFC activity may be more directly related to performance optimization than the corresponding left PFC region. Potential mechanisms include dynamic control of attention or listening strategy, streamlining working memory or processing efficiency, among many others. Some subjects reported that they actively explored different listening strategies during the dichotic task blocks, particularly regarding which ear to focus on or ignore. Exploratory spatial listening adjustments involved in right/left attention switching may thus have contributed to the CBF signals. Indeed, the DLPFC receives from auditory cortex afferent information streams related to spatial localization of sounds (Romanski et al., 1999). Future experiments in which subjects are explicitly instructed to alternate attention between the ears could more specifically guide this line of inquiry.

The EEG findings of enhanced gamma power at parietal and frontal locations during dichotic listening tasks is consistent with previous work showing that gamma spectral power is correlated with working memory engagement in both auditory and visual sensory modalities (Tallon-Baudry et al., 1998; Lutzenberger et al., 2002; Kaiser et al., 2003). The slight, albeit not statistically significant, reduction in EEG gamma power during dichotic block 2 may reflect the fact that some of the reduced subjective effort was associated with a more streamlined use of working memory. Unsurprisingly, all subjects agreed the dichotic tests were significantly more difficult than diotic, a subjective report that was supported quantitatively by the heartrate observations (Fig. 2C-D) as well as the greater regional CBF activation during dichotic listening. This result is consistent with fMRI measurements under both diotic and dichotic listening tasks (Thomsen et al., 2004) and is expected given that diotic listening does not involve a significant memory recall burden.

Importantly, we were able to use optical techniques to reproduce dichotic listening findings that were previously obtained only through fMRI. The average noise level perceived by subjects was minimal, particularly given the added hearing protection from the foam coupler tips of the insert earphones. Functional hyperemia dynamics, both onset and offset, were fast; after the final digit combination was delivered in any dichotic or diotic listening block, CBF returned to rest levels on a timescale of ∼15 sec. Our ongoing measurements of regional CBF exhibited a gradual upward change over the ∼30-minute course of recording likely owing to subject fatigue, an issue common to most block-paradigm behavioral experiments (Petersen and Dubis, 2012).

The incidental DPOAE findings are provocative in that they invoke relevance of infrastructure for longer-range, top-down listening optimization via olivocochlear efferents. Medial olivocochlear efferents, which can be modulated by activity in the auditory cortex (Perrot et al., 2006), modulate the nonlinear mechanical properties of the inner ear by altering outer hair cell stiffness through electromotility, a process mediated by the voltage-sensitive motor protein prestin (Liberman et al., 2002; Dallos et al., 2006). Andeol et al. found that sound localization tasks in humans elicited alterations in evoked otoacoustic emissions, a proxy for cochlear mechanical state (Andéol et al., 2011b). Wittekindt et al. found that attention-modulated changes in DPOAEs during behavioral tasks are associated with concomitant changes in EEG (Wittekindt et al., 2014). Subjects who displayed the largest relative change in regional cerebral blood flow in PFC also had the largest baseline left year DPOAEs; do these subjects utilize corticofugal efferent control of the cochlea to a greater extent than others? Are these subjects more “dexterous” in their selective listening ability or capacity for optimizing performance? Does the finding merely represent an unrelated epiphenomenon? Future studies that insert periodic measurements of DPOAEs during an experiment could help assess potential roles for corticofugal efferent control—and the relationship with PFC regions therein—during dichotic listening.

## Conclusions

Current hypotheses regarding the role of the right PFC during dichotic listening focus on (1) compensation for the left hemisphere speech and language advantage, (2) a putatively distinct, yet still speech/language related role, or (3) task-neutral, performance optimization roles such as managing working memory, attention, or other aspects of information professing. While our findings cannot categorically confirm or reject any of these classes of proposed mechanisms, our results do support the hypothesis that the right DLPFC is more involved in performance optimization than left during repeated dichotic listening tasks. Future measurements with the same apparatus yet different sensory modalities may help elucidate the degree to which this asymmetry is behavioral task- or sensory modality-neutral.

## Acknowledgements

We thank Drs. R. Jacobson, W. Silver, J. Mast, P. Overby, and S. Parvez for help with subject recruitment and for helpful conversations. We thank Drs. W. Baker and A. Parthasarathy for technical assistance with the optical apparatus. We additionally thank Dr. H. Jang for assistance developing the experimental protocol. This work was supported by recruitment funds from New York Medical College (JANF) and startup funds from the Margaret Milam McDermott Distinguished Chair in Anesthesiology and Pain Management at the University of Texas Southwestern Medical Center (DRB).

